# Exploring the impact of social relevance on the cortical tracking of speech: viability and temporal response characterisation

**DOI:** 10.1101/2025.09.23.674728

**Authors:** Emily Y.J. Ip, Asena Akkaya, Martin M. Winchester, Sonia J. Bishop, Benjamin R. Cowan, Giovanni M. Di Liberto

**Affiliations:** School of Computer Science and Statistics, University of Dublin, Trinity College, Dublin 2, Ireland; ADAPT Centre; Trinity College Institute of Neuroscience, University of Dublin, Trinity College, Dublin 2, Ireland; School of Psychology, University of Dublin, Trinity College, Dublin 2, Irelan; School of Information and Communication Studies, University College Dublin, Dublin 4, Ireland

## Abstract

Human speech is inherently social. Yet our understanding of the neural substrates underlying continuous speech perception relies largely on neural responses to monologues, leaving substantial uncertainty about how social interactions shape the neural encoding of speech. Here, we bridge this gap by studying how EEG responses to speech change when the input includes a social element. In Experiment 1, we compared the neural encoding of synthesised undirected monologues, directed monologues, and dialogues. In Experiment 2, we extended this by using podcasts, addressing the additional challenges of real speech dialogue, such as dysfluency. Using temporal response function analyses, we show that the presence of a social component strengthens the cortical tracking of the speech envelope, despite identical acoustic properties. Neural responses to synthesised speech showed a strong correlation with those for real speech podcasts, with a stronger alignment emerging for more socially-relevant speech material. In addition, we demonstrate that robust neural indices of sound and lexical-level processing can be derived using real podcast recordings despite the presence of dysfluencies. Finally, we present a simulation to put to the test the robustness of temporal response function analyses under increasing levels of dysfluency. Together, these findings highlighting the impact of social elements in shaping auditory neural processing, providing a framework for future investigation and analysis of social speech listening and speech interaction.

**Significance Statement:** Human speech is rarely produced or processed in a social vacuum. Yet, our understanding of continuous speech neurophysiology mostly comes from experiments involving speech monologues. This study reveals how social context modulates the neural encoding of speech. We directly contrast neural signals recorded when participants listened to monologues and dialogues, using controlled material from speech synthesis and real podcast recordings. We found that the social element amplifies the neural encoding of speech features, reflecting greater engagement. We also show strong correlation between synthetic and real podcast neural responses, scaling with social relevance. Finally, we demonstrate that lexical processing can be measured robustly even amid natural dysfluencies. These insights advance our understanding of speech neurophysiology, informing future research on social speech.

## Introduction

To understand speech, our brains must rapidly process sounds into linguistic meaning (Friederici, 2011; Poeppel et al., 2012). The neural basis of this phenomenon has been studied extensively using controlled paradigms involving targeted manipulations (Carreiras et al., 2005; Kutas & Federmeier, 2011; Osterhout & Holcomb, 1993), and more recently with listening tasks involving more realistic continuous streams of speech (Di Liberto & Ip, 2025; Di Liberto et al., 2021; Gwilliams et al., 2023). That work utilised a variety of neural recording technologies, including electro- and magneto-encephalography (EEG and MEG) (Défossez et al., 2023; Destoky et al., 2019), functional magnetic resonance imaging and functional near-infrared spectroscopy (Butler et al., 2020; Chen et al., 2020; Formisano et al., 2008), and intracranial electrical recordings (e.g., electrocorticography and stereotactic EEG) (Bernabei et al., 2021; Hamilton et al., 2021; Mesgarani et al., 2014). Overall, this research has led to a new understanding of the sound-to-meaning transformation, providing detailed insights into the spatial and temporal dynamics that unfold throughout that process (Kocagoncu et al., 2017; Uusvuori et al., 2008). However, the necessary experimental constraints have led to listening tasks centred around audiobook-style, pre-recorded speech material, while ignoring one of the key functions of speech: the social element.

Recent research has explored speech interaction, for example by measuring brain-to-brain synchrony using simultaneous neural recordings from interacting individuals (Kinreich et al., 2017; Schwartz et al., 2024). However, those interactive experiments are far removed from previous speech listening research, as other factors at play (e.g., social anxiety, familiarity, turn-taking) make it difficult to relate findings across the two domains. To achieve this and advance the study of social speech neurophysiology, we require a mechanistic understanding of how specific social factors impact the neural processing of speech, which requires careful experimental control. The present study takes a step in that direction, by extending the conventional audiobook-style listening paradigm to stimuli with progressively higher social relevance. In doing so, we investigate how challenges of real speech (e.g., dysfluencies) impact our ability to measure the neural processing of speech and language (Di Liberto & Ip, 2025; Mousavi et al., 2024).

We present results from two EEG experiments. The first experiment compares the neural encoding of speech while participants listened to pre-recorded monologues and dialogues (**Experiment 1**). *Dialogue* typically involves turn-taking, mutual feedback, and shared communicative goals (Garrod & Pickering, 2004), while monologues lack immediate (auditory) reciprocity. Nevertheless, monologues can also include a social dimension by being *directed* to a listener, as opposed to *undirected monologues*, when they are not aimed at any particular listener. Using the temporal response function analysis (TRF), this study tests if social relevance impacts the neural processing of speech during listening. This leads to our first hypothesis:

### Hypothesis 1

Social relevance enhances the cortical encoding of speech, leading to stronger encoding for directed than undirected monologues.

Synthesised speech was used to control for acoustic and lexical differences across conditions, ensuring that differences in the EEG would be due to social relevance as opposed to other factors such as prosody, word rate, or volume level (see **Methods**). To generalise the results from Experiment 1 to the case of real human speech, we also conducted a second experiment using real dialogue recordings from popular podcasts (**Experiment 2**). The material included speech from different registers to capture a broader range of speech types, increasing generalisability. Furthermore, real spontaneous includes dysfluency(Shriberg, 2001), which can challenge natural speech comprehension, for example by hampering syntactic parsing (Ferreira & Bailey, 2004), contributing to processing delays that impact word predictability (Corley et al., 2007). Here, we investigate the possibility of studying speech material including dysfluencies, testing if robust EEG measurements of speech processing can be derived at varying levels of dysfluency, with the following hypotheses:

### Hypothesis 2

The social element of speech leads to a consistent neural signature, which generalises across synthesised and real speech dialogue.

### Hypothesis 3

Neural measurements of speech encoding are robust to dysfluency in the speech material.

## Materials and Methods

### EEG Experiment 1

#### Data acquisition and experimental setup

Twenty healthy individuals (10 male, 10 female; between 18 and 30 years old, median = 22) participated in the experiment. All participants were native English speakers and reported no history of hearing impairment or neurologic disorder, provided written informed consent, and were paid for participating. The study was undertaken in accordance with the Declaration of Helsinki and was approved by the ethics committee of the School of Computer Science and Statistics at Trinity College Dublin. The experiment was carried out in a single session for each participant. EEG data were recorded from 64 electrode positions, digitised at 512 Hz using a BioSemi ActiveTwo system. Audio stimuli were presented at a sampling rate of 48 kHz using Sennheiser HD 650 headphones and the Psychtoolbox-3 MATLAB library (Brainard, 1997). Testing was conducted at Trinity College Dublin in a dark room. Participants were instructed to maintain visual fixation on a crosshair centred on the screen and to minimise motor activities during the speech listening task.

#### Stimuli

Monologue scripts were generated with the large language model (LLM) GPT-4 (OpenAI, 2024), using prompts such as “Could you create a monologue, with a duration of approximately 2 minutes?”. From 30 generated monologues, scripts with highly similar content and those containing strongly emotional themes (e.g., scenarios involving extreme sadness or anger) were excluded to avoid potential confounds in the experiment. After this filtering process, 16 monologue scripts with sufficiently varied topics were retained for use as stimuli. The selected monologues were further processed with GPT-4, generating eight directed and eight undirected monologues. That manipulation was achieved by using prompts such as: *“Turn this into a directed monologue that addresses the listener”* and “*Turn this into an undirected monologue, where the speaker talks to themselves.”*

The 16 monologues (8 directed, 8 undirected) were then converted into dialogues using the same approach, with the prompt: “*Convert the following monologue into a dialogue between two speakers while preserving the original content. Keep the duration approximately the same and distribute the information naturally across the speakers.*” The aim of this transformation was to create dialogue versions that preserved the semantic content of the original monologues while introducing an interlocutor. This approach allowed us to maintain comparable content across monologue and dialogue conditions, thereby minimising potential confounds related to topic or narrative structure.

Scripts were synthesised using the realistic AI-based voice generation software PlayHT version 2.0^1^. The generated speech sounds were manually inspected sentence-by-sentence to minimise generation artifacts. Speech segments with clear artifacts or sounding unnatural were replaced by re-running the generation for the affected segment, ensuring a consistent tone and emphasis. Note that this was possible as each generation produces a different result. Altogether, this procedure generated 32 sound files (16 monologues, 16 dialogues) synthesised using different voices (16 voices: 8 male, 8 female). The final version of auditory stimuli has been deposited in the OSF repository^2^.

To control for potential acoustic and speech-structure confounds, several acoustic properties of each stimulus, including duration, RMS amplitude, mean fundamental frequency (F0), F0 variability (standard deviation), and word rate (number of words per second) were quantified. Participants (N=15) participated in an online survey, where they were asked to categorise whether speech recordings were undirected or directed. Participants were presented with 16 speech recordings, each ∼2-min of duration. Acoustic measures were compared across conditions (undirected-directed monologues, dialogues) using Kruskal-Wallis tests. No significant differences were observed for any of the acoustic properties across the three conditions (all Kruskal-Wallis tests, *p* > 0.05; see **Table S3**).

In addition, monologue stimuli (combination of directed and undirected conditions) were compared with dialogue stimuli, showing no statistically significant differences for any of the acoustic properties (all Wilcoxon rank-sum tests, *p* > 0.05; see **Table S3**).

#### Experimental procedure

All thirty-two stimuli were presented in a single experimental session in a randomised order, which was different across participants. Participants were free to take breaks of arbitrary length in-between stimuli. To maintain engagement throughout the experiment, participants were periodically (25% of the trials) asked to rate how interesting they found the speech segments on a 5-point scale (1: not at all interesting; 3: neutral; 5: very interesting).

### EEG Experiment 2

#### Data acquisition and experimental setup

Twenty healthy individuals (10 male, 10 female; between 20 and 39 years old, median = 26), separate from Experiment 1, participated in the second EEG experiment. All participants were native English speakers and reported no history of hearing impairment or neurologic disorder, provided written informed consent, and were paid for participating. The study was undertaken in accordance with the Declaration of Helsinki and was approved by the ethics committee of the School of Psychology at Trinity College Dublin.

The experiment was carried out in a single session for each participant. EEG data were recorded from 64 electrode positions, digitised at 512 Hz using a BioSemi ActiveTwo system. Audio stimuli were presented at a sampling rate of 44,100 Hz using Sennheiser HD 280 Pro headphones and the PsychoPy Python library version 2022.2.5 (Peirce et al., 2019). Testing was conducted at Trinity College Dublin in a dark room. Participants were instructed to maintain visual fixation on a crosshair centred on the screen and minimise motor activities during the speech listening task.

#### Stimuli

Podcast recordings were selected from publicly available sources: 1) American podcast shows called *Brains On!* and *Forever Ago;* 2) *WIRED 5 Levels of Difficulty* series on YouTube; and 3) interviews with the same two hosts *Forever Ago,* available online^3^. Dialogues contained general interest topics, such as questions about science and math (e.g., why do we sweat, why memories as a young child are forgotten). Altogether, forty-two podcast recordings involving 1-to-1 dialogues were selected for this experiment (the same stimuli for all participants), totalling one hour in duration. Sixteen of those recordings involved adult-adult (AA) dialogues, while the remaining twenty-six were adult-child (AC) dialogues. The AA and AC dialogues contained the same five adult speakers (podcast hosts), speaking to a range of different children and other adults. The higher number of recordings for the latter condition ensured that the two conditions had a comparable number of words, as child-speech and child-directed speech are generally slower than adult-speech (Ferreira, 2019; Snow, 1972). To that end, the selected speech registers captured a range of speaking styles for testing the robustness of our findings (e.g., the impact of dysfluencies on the EEG encoding of speech) to variations in acoustic and lexical properties of a speech input.

#### Experimental procedure

Participants were presented with the forty-two speech stimuli (referred hereafter as trials) in a random order, separated by a short break. To encourage attention, participants were asked to rate how interesting they found the speech segment on a scale from 1-5 (1: not at all interesting; 3: neutral; 5: very interesting) after each trial. Additionally, for a random subset of twenty percent of the trials, the participants were asked a behavioural multiple-choice question to select the set of keywords that best described the trial they just heard. This was also to keep the participants engaged and on task throughout the experiment.

### EEG data preprocessing

Neural data from both experiments were analysed offline using MATLAB software (MathWorks), using the same preprocessing and analysis pipeline. EEG signals were digitally filtered using Butterworth zero-phase filters with order 2, implemented with the function *filtfilt*. The primary analyses were carried out on EEG data band-pass filtered between 0.5 and 8 Hz, as that range has been shown to be particularly sensitive to the cortical tracking of speech (Di Liberto et al., 2015; Doelling et al., 2014; O’Sullivan et al., 2015; Vanthornhout et al., 2018). Low-frequencies from 0.5 Hz to 1 Hz were included as they were previously shown to be important for measuring the neural encoding of lexical predictions with EEG (Broderick et al., 2018; Carta et al., 2024). Secondary analyses were carried out to pinpoint specific sub-bands driving effects of interest. In Experiment 1, we considered the delta-band (0.5-4 Hz), theta-band (4-8 Hz), alpha-band power (8-12 Hz), beta-band power (12-30 Hz) and broadband (0.5-30 Hz). In Experiment 2, we considered just the delta-band (0.5-4 Hz) and theta-band (4-8 Hz), as they led to the highest EEG prediction correlation in Experiment 1 (**Figure S1**). The power of alpha and beta bands was calculated as the instantaneous magnitude of the Hilbert transform in those sub-bands. This selection is based on evidence that alpha and beta rates reflect predictive processing, attention, and working memory (Haegens et al., 2017; Samaha et al., 2015). EEG signals were down sampled to 64 Hz, reducing computational and storage costs. EEG channels with a variance exceeding three times that of the surrounding ones were replaced by a spherical spline interpolation (Delorme & Makeig, 2004; Perrin et al., 1989). Signals were then re-referenced to the average of the two mastoid channels.

### Stimulus features

The present study examines the neural encoding of speech and language features during speech listening. The same features and analyses were used across the two experiments.

#### Sound envelope and derivative

The sound envelope was calculated as the instantaneous magnitude of the Hilbert transform of the sound-pressure waveform. The speech envelope captures the slow amplitude modulations of the speech signal, which have been shown to be robustly reflected in the EEG signal (Crosse et al., 2016a; Ding & Simon, 2012; Lalor & Foxe, 2010). The envelope-related features assessed in this study comprised the sound envelope and half-way rectified envelope derivative (calculated from the envelope at the original sampling rate with the MATLAB *diff* function, and by replacing negative values with zero). Matching the processed EEG signal, these acoustic features were then down sampled to 64 Hz.

#### Word onsets

The forced alignment software DARLA (Reddy & Stanford, 2015) was used to produce a word-by-word time-alignment between each speech segment and its transcript. The onset time of all words in the speech stimuli was then coded into sparse vectors of zeros and ones with the same length and sampling rate as the other features, where ones represent the onset time of a word, consistent with previous studies (Broderick et al., 2018; Chalehchaleh et al., 2025; Di Liberto et al., 2021).

#### Lexical surprise and entropy

The second experiment in the present study investigates the possibility of measuring neural signatures of lexical surprise and entropy in the presence of dysfluencies, which are typically part of dialogues (Bosker et al., 2014; Clark & Tree, 2002; Fox Tree, 2001; Tree, 1995). Next-word lexical surprise and entropy values were extracted from Mistral-7B-Instruct-v0.1 (Jiang et al., 2023), an open source state-of-the-art LLM that has been fine-tuned on conversation datasets. For each trial, lexical surprise was calculated as the negative log contextual probability of each word (Heilbron et al., 2022; Slaats & Martin, 2025). Additionally, for each trial lexical entropy was calculated as the weighted sum over the surprise values for all possible next words, and measures the level of uncertainty the model has for its next-word prediction (Goldstein et al., 2022; Slaats & Martin, 2025). Lexical surprise and entropy feature vectors were built by scaling the word onset vector by its corresponding surprise and entropy value, respectively. Previous work employing similar models (e.g., GPT-2) found lexical expectation values to reliably explain EEG/MEG variance when considering speech listening tasks (Dou et al., 2024; Heilbron et al., 2022). However, that work typically involved scripted speech monologues without dysfluencies.

#### Dysfluencies

To investigate the impact of dysfluencies on lexical surprise and entropy and their neural signatures, we built two lexical surprise and entropy vectors: (1) one that included dysfluencies and (2) one that excluded dysfluencies in the speech material (**Figure 4A**). Such dysfluencies were manually transcribed, annotated, and removed (see the dysfluency types considered in **Table S1**). To balance the amount of data in both versions of the surprise and entropy vectors, the annotated dysfluencies were removed from both feature vectors. Therefore, the main difference in the two versions of the surprise and entropy vectors lies primarily in if/how the presence of dysfluencies impacts the predictions of upcoming words. In practice, this led to two surprise and entropy feature vectors with the same word onsets, but different surprise and entropy values.

### Temporal response function analysis

A system identification approach was employed to determine how specific stimulus features map onto EEG activity. This approach, known as the Temporal Response Function (TRF) (Ding & Simon, 2012; Edmund C. Lalor, 2009), utilises regularised ridge linear regression (Crosse et al., 2016) to estimate a filter, or model, that best captures how the brain converts input stimulus features into the resulting neural responses. One of the two metrics derived from the TRF is the weights of the model over time. The brain’s response to a speech stimulus is not instantaneous; rather, it unfolds over time when speech is heard. Therefore, the brain’s electrical activity is affected not just at the moment the sound occurs, but also over a subsequent time period. This delayed and extended influence is captured in the time lags of the TRF weights between the stimulus and the EEG response. These time lags can be analysed both spatially (across the scalp) and temporally (in terms of response delays).

As is typical (Crosse et al., 2016b; Crosse et al., 2021), to evaluate the model’s ability to generalise and avoid overfitting, we used leave-one-out cross-validation across trials. In this process, TRF models were trained on all but one trial and then used to predict the EEG signal for the left-out trial. The TRF model performance is then assessed by computing Pearson’s correlation between the predicted EEG signal and the actual EEG signal on the left-out fold at each scalp electrode. This constitutes the second metric derived from the TRF: the prediction correlation, or the EEG variance explained by a specific set of speech feature(s). The EEG variance captured by speech features is interpreted as how well the model was able to map the encoding of speech features in the brain signals. Fitting the model with one speech feature results in a *univariate model,* while fitting the model while more than one speech feature results in a *multivariate* model.

Using these objective metrics, univariate forward TRF models were used to assess the corresponding neural response to each speech feature of interest. Three univariate TRF models were fit in Experiments 1 and 2 for the following speech features: the speech envelope, envelope derivative, and the word onset. In Experiment 1, the strength of the neural response to the speech in each of the three conditions was assessed through the TRF models’ prediction correlations for any effect of social relevance. In Experiment 2, the temporal dynamics of the EEG response to podcast dialogues were examined from the TRF weights, expecting latencies consistent with previous results on audiobook listening tasks. Despite variability in the acoustics of the spontaneous dialogue stimuli across speech registers (Ferreira & Ferreira, 2019; Snow, 1972) and, as a result, the neural responses, the TRF measured is designed to remain robust and stable against such specific differences as it estimates the consistent impulse response of the brain. This serves as an opportunity to build speech dialogue TRFs that are stable across various types of speech and speech registers in real dialogues. Notably, we expected to measure a negative deflection of the TRF at latencies around 400 ms (i.e., the TRF-N400 response) for this podcast listening task that, compatibly with previous monologue listening work (Broderick et al., 2018; Broderick et al., 2021; Heilbron et al., 2022), would index word-level processing.

Multivariate TRF (mTRF) analyses were also conducted in Experiment 2 to assess if the cortical encoding of lexical surprise and entropy was more aligned with a next-word prediction model including or disregarding dysfluencies. This enables us to evaluate how much, if at all, dysfluencies impact lexical predictions during speech, informing us on the most appropriate approach for measuring lexical expectations in the human brain in contexts involving real speech (i.e., with dysfluencies) (Heilbron et al., 2022; Mousavi et al., 2024). Sound envelope and word segmentation information features (e.g., speech, speech derivative, word onset) were concatenated to represent the stimulus, along with one of two versions of surprise: either dysfluent or non-dysfluent surprise.

### Statistical analyses

Parallel statistical procedures were applied to the datasets from the two experiments. Statistical analyses were performed on EEG prediction-correlation values obtained from the TRF models. For each participant and condition, prediction correlations were averaged across all EEG electrodes to obtain subject-level values, and these values were used in all statistical tests. To determine whether the TRF models captured meaningful neural tracking of the speech features, prediction correlations were tested against zero using one-sample *t*-tests.

To compare experimental conditions, repeated-measures statistical tests were used because all conditions were recorded from the same participants. When three levels of social relevance (undirected monologue, directed monologue, and dialogue) were compared, a one-way repeated-measures ANOVA was applied to the subject-level averages. When the omnibus test was significant, pairwise comparisons between conditions were performed using Wilcoxon signed-rank tests.

For analyses of TRF weights across time lags, multiple comparisons were controlled using the false discovery rate (FDR) procedure with *p* = 0.05, according to the procedure set out by (Benjamini & Yekutieli, 2001) for dependence assumptions.

A cross-experiment comparison was carried out by correlating TRFs for the two datasets. Considering that the two datasets differ in both stimuli and participants, we carried out that comparison via a bootstrap resampling procedure on the group-averaged TRF weights. Pearson’s correlation scores were calculated for the TRF weights corresponding to univariate envelope, envelope derivative, and word onset features. The general assumption in averaging TRFs (as for ERPs) is that TRFs for individual participants may vary, but that participants share certain consistent dynamics, which would emerge by averaging. Here, the assumption was that averaging TRFs from two distinct samples of participants (from the same population) would lead to the same TRF (when enough participants are available), with higher TRF correlation for more similar experimental tasks. Accordingly, we expected higher inter-experiments TRF correlations when considering more similar tasks (i.e., dialogue listening). This test was carried out via a bootstrap resampling procedure that derived 1,000 correlations between the average TRFs for each of the conditions in Experiment 1 and for Experiment 2, generated using a random sample with replacement on each participant cohort. Because Mauchly’s test indicated that the assumption of sphericity was violated (*p* < 0.05), a nonparametric Friedman test was conducted to examine differences in the correlations across conditions for each TRF feature using JASP software (Love et al., 2019).

The dysfluency analysis in Experiment 2 included repeated measures ANOVAs. Post-hoc tests using paired *t-*tests with the Holm correction were performed to examine the effect of including/excluding dysfluencies on TRF model predictions, from which we report the adjusted *p-*value. Matlab was used for statistical analysis in Experiment 1. For Experiment 2, statistical analyses were performed using Matlab and JASP software.

### Code and data accessibility

The TRF analysis was conducted using the publicly available and open source mTRF-toolbox version 2.6 (Crosse et al., 2016a; Crosse et al., 2021), which can be downloaded from https://github.com/mickcrosse/mTRF-Toolbox or https://sourceforge.net/projects/aespa/. Stimulus features and EEG data were stored according to the continuous-event neural data structure CND (Di Liberto et al., 2024). Analyses were conducted using scripts shared via the GitHub channel^4^ of the Cognition and Natural Sensory Processing (CNSP) and Natural Sensory Processing open science initiative^5^. Custom scripts were used for data visualisation and statistical analyses. Datasets for Experiments 1 and 2 will be publicly shared via the open science repository OSF (osf.io) at the time of publication and will also be included in the list of publicly available datasets of the CNSP-initiative.

## Results

### Experiment 1

The survey indicated that participants could categorise undirected versus directed speech sounds, achieving an accuracy of 0.94 (average across participants; ± 0.11 SD). Performance was significantly above chance level, as confirmed by one-sample t-tests (overall: *t*(14) = 15.9, *p* = 1.1 × 10⁻³²; undirected stimuli: *t*(14) = 13.2, *p* = 3.1 × 10⁻²¹; directed stimuli: *t*(14) = 17.4, *p* = 2.4 × 10⁻³³; **Fig. S4**). Monologues were rated as more interesting on average than dialogues (M_monologue_ = 3.42, M_dialogue_ = 2.87; paired *t*-test; *t*(18) = −2.26, *p* = 0.036), while no statistically significant difference in interest ratings emerged between undirected and directed monologues (M_undirected_ = 3.72; M_directed_ = 2.89; paired *t*-test: *t*(8) = −2.13, *p* = 0.066).

Univariate TRFs were fit to model the relationship between stimulus features (speech envelope, envelope derivative, and word onset) and the broadband EEG signal (0.5-30 Hz). Separate TRF models were fit for the three stimulus features. First, we report results for the widely used sound envelope feature as in previous studies (Muncke et al., 2022; Vanthornhout et al., 2018). EEG prediction correlation values calculated with leave-one-out cross validation (see **Materials and Methods**) had magnitudes compatible with envelope TRF analyses in previous research (r∼0.2-0.4, after averaging across all participants and EEG channels). Comparing EEG prediction correlations for envelope TRFs across conditions (undirected monologue, directed monologue, and dialogue) revealed a statistically significant effect of social relevance (one-way repeated measures ANOVA, *F*(2, 38) = 5.98, *p* = 0.006), with more socially-relevant stimuli producing larger EEG prediction correlation values. Post-hoc analyses indicated lower prediction correlations in the undirected monologue condition compared with both the directed monologue (Wilcoxon signed-rank tests, *p* = 0.008) and dialogue conditions (*p* = 0.003; **Figure 1A**). No statistically significant differences were observed between directed monologue and dialogue conditions *(p* = 0.89). No statistically significant effects of social relevance were observed for the other stimulus features considering broadband EEG (envelope derivative: *F*(2, 38) = 2.16, *p* = 0.129; word onset: *F*(2, 38) = 1.03, *p* = 0.366).

**Figure 1.**
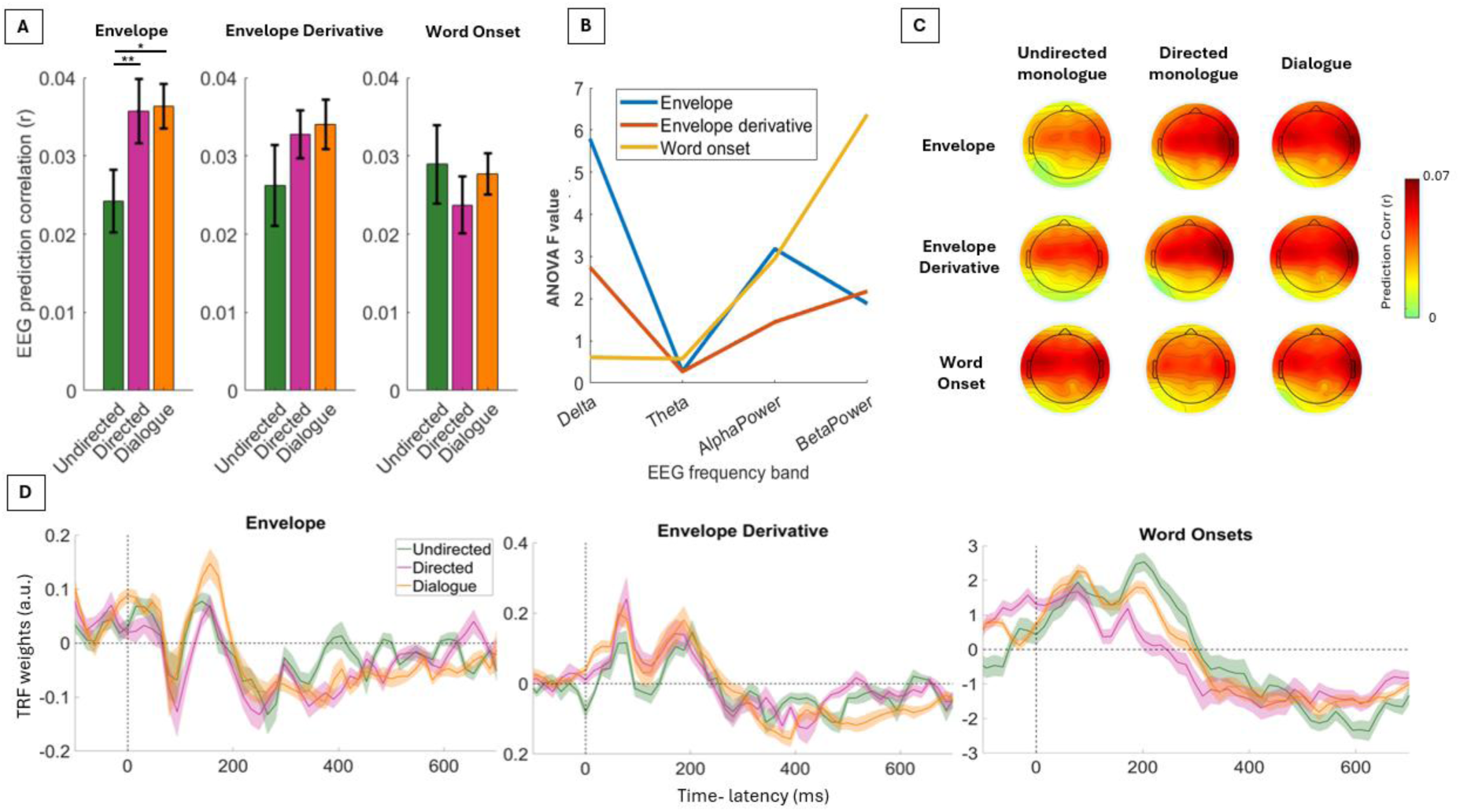
Social relevance impacts the cortical tracking of speech. **(A)** Univariate forward TRFs were fitted for envelope, envelope derivative, and word onset features to predict the on the broadband EEG in Experiment 1. EEG prediction correlations (Pearson’s r) were first averaged across electrodes within each participant, and group means (± SEM across participants) are shown for the three experimental conditions: undirected monologue, directed monologue, and dialogue.Envelope TRFs for the socially-relevant conditions exhibited stronger EEG prediction correlations (one-way repeated measures ANOVA with Wilcoxon signed-rank post hoc comparisons; *p < 0.05, **p < 0.01). **(B)** The same analysis was repeated for all EEG sub-bands of interest. F-value of the repeated measures ANOVA is reported for each sub-band, indicating the effect of social relevance across features and EEG frequency bands. The strongest effects of social relevance for the envelope TRF emerged in the delta-band, and for word-onset in the gamma-band. **(C)** Scalp topographies of the EEG prediction correlations for each condition and speech feature. Warmer colours indicate higher prediction correlation values. All topographies showed qualitatively similar spatial patterns, compatible with previous studies on continuous speech listening (Di Liberto et al., 2015). **(D)** TRF weights averaged across all participants at channel Cz are shown for the three conditions and stimulus features. Shaded areas represent the standard error of the mean (SEM) across participants.

TRF and ANOVA analyses were repeated for different EEG sub-bands to determine if the effect was specific to certain cortical rhythms. The envelope TRF analysis showed strongest effects in low-frequency EEG (**Figure 1B**), with a statistically significant effect in the delta-band (F(2, 38) = 5.80, p = 0.006); a trending effect in the alpha-band (F(2, 38) = 3.18, p = 0.053); and no statistically significant effects elsewhere (theta-band: *F*(2, 38) = 0.27, *p* = 0.763; beta-band: *F*(2, 38) = 1.89, *p* = 0.166). Conversely, the word onset TRF showed strongest effects in higher EEG frequencies, with a statistically significant effect in the beta-band (one-sample ANOVA, *F*(2, 38) = 6.38, *p* = 0.004); a trending effect in the alpha-band (*F*(2, 38) = 2.95, *p* = 0.064); and no statistically significant effects elsewhere (delta-band: *F*(2, 38) = 0.61, *p* = 0.551; theta-band: *F*(2, 38) = 0.57, *p* = 0.559). No statistically significant effects were measured for the envelope derivative TRF (delta-band: *F*(2, 38) = 2.75, *p* = 0.077; theta-band: *F*(2, 38) = 0.27, *p* = 0.765; alpha-band: *F*(2, 38) = 1.45, *p* = 0.248; gamma-band: *F*(2, 38) = 2.17, *p* = 0.128).

We also report TRF weights for the models in the EEG broadband. Based on previous research, we expected a P1-N1-P2 pattern with latencies around 40, 80, and 140ms respectively (Crosse et al., 2016a; Di Liberto et al., 2015; Lalor & Foxe, 2010), where longer latencies have been linked with factors such as selective attention (Power et al., 2012). The literature also informs us on the neural dynamics corresponding with higher order features like word onsets, such as a large slow negative deflection at a latency of approximately 400 ms (Broderick, 2018). Here, we found TRF patterns that were qualitatively similar to that literature in all those aspects.

### Experiment 2

EEG signals were recorded as participants listened to spontaneous two-party dialogues from popular podcasts, each involving a podcast host and a guest. Forward univariate TRFs were derived for the two speech features showing an effect of social relevance: the sound envelope and word onset features (**Figure 2**; we also report the envelope derivative results in **Figure S2**). As in Experiment 1, EEG prediction correlations were calculated with leave-one-out cross validation for each stimulus feature separately. EEG prediction correlations were above chance for both sound envelope (**Figure 2A-left**; *t*-tests, *p* < 0.001) and word onsets (**Figure 2B-left**; *t*-tests, *p* < 0.001), reflecting a statistically significant EEG encoding of the two features.

**Figure 2.**
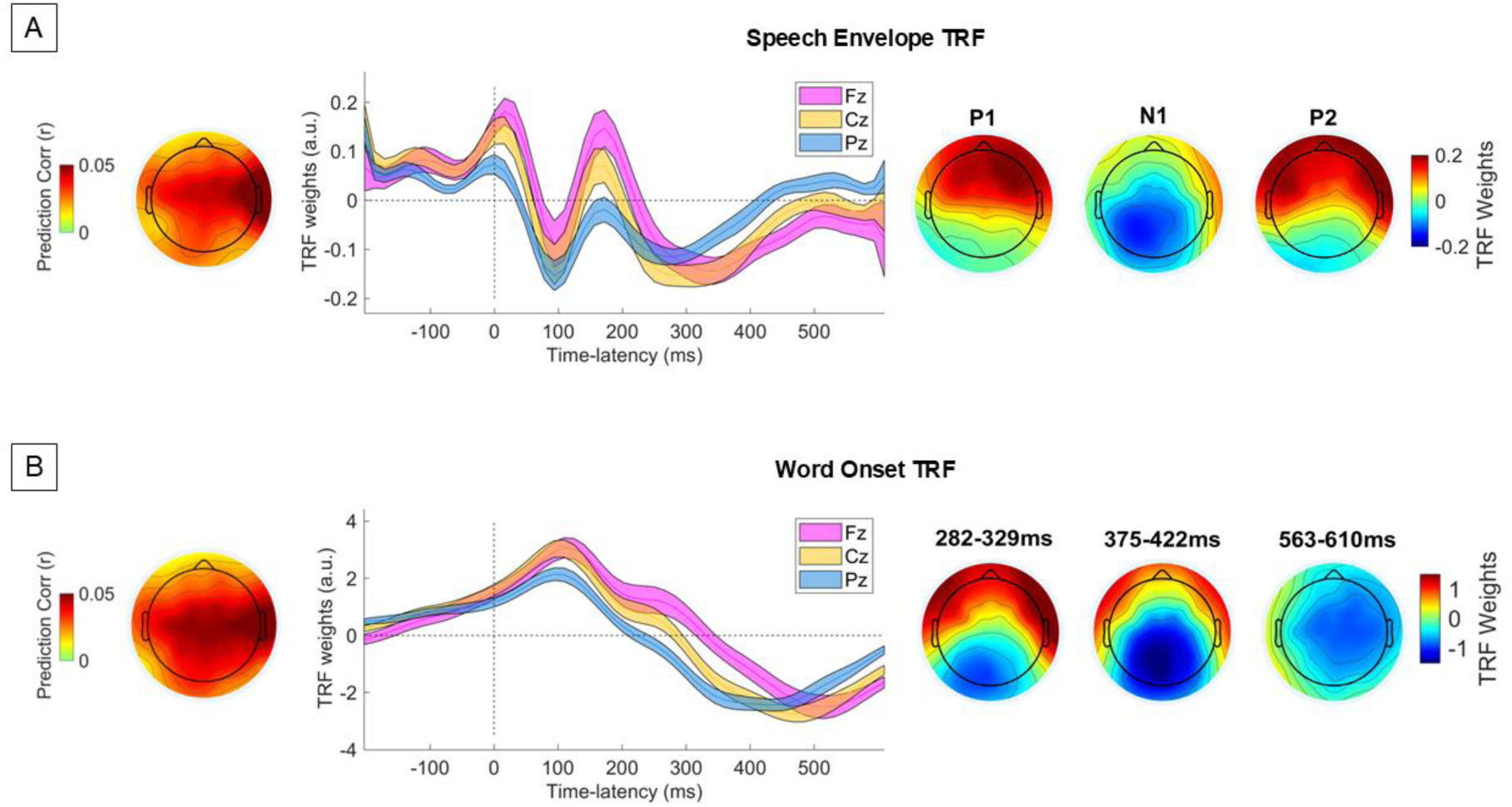
Characterising the cortical tracking of speech using real podcast dialogues. Univariate forward TRF models in Experiment 2 for envelope **(A)** and word onset **(B)** features. These analyses aggregated data from both listening conditions (AA and AC). Scalp distributions of the average EEG prediction correlations are reported (left panels). TRF weights at channels Fz, Cz and Pz (middle-panels) and the TRF weights at three key time-lags exhibit patterns compatible with the literature (Chalehchaleh et al., 2025; Di Liberto et al., 2015; Lalor, 2009).

In Experiment 2, the podcast host was always an adult speaker, while the podcast guest was either an adult or a child. This led to two conditions (adult-adult conversations - AA; adult-child conversations - AC) that we compared to determine if their multi-faceted differences (e.g., prosody, lexis, word-rate) would impact the envelope and word onset TRF model fits. While there may be other differences that exist elsewhere between AA and AC speech, for the purposes of the current analyses, we focused on just these two speech features. EEG prediction correlations averaged across scalp locations did not differ significantly between AA and AC stimuli (Wilcoxon paired signed-rank test (\**p* <0.05): envelope: *p* = 0.177; word onset: *p* = 0.165). The effect of stimulus type (AA versus AC) upon TRF feature weights at channel Cz was also examined for each time lag. No statistically significant differences emerged (FDR-corrected Wilcoxon paired signed-rank test, \**p* > 0.05 across all time lags). On this basis, the analyses that follow are carried out by combining AA and AC stimuli. While no effects of speech register emerged in this investigation, further work could investigate possible effects on other linguistic features (Piazza et al., 2025).

The TRF weights were qualitatively comparable with the previous literature (Di Liberto et al., 2015; Lalor et al., 2009) and with the results from Experiment 1, with the envelope TRF presenting the typical P1-N1-P2 characteristics, and the word onset TRF exhibiting the typical dominant negative deflection at about 400 ms (**Figure 2-middle**). We measured Pearson’s correlations between the group TRF weights across the two experiments. The resulting group-level correlations after averaging the TRF weights of the 64 EEG channels across participants are shown in **Figure 3**, indicating high correlations (up to 0.92) across experiments, with average correlation values increasing with social relevant i.e., undirected monologue, directed monologue, synthesised dialogue.

**Figure 3.**
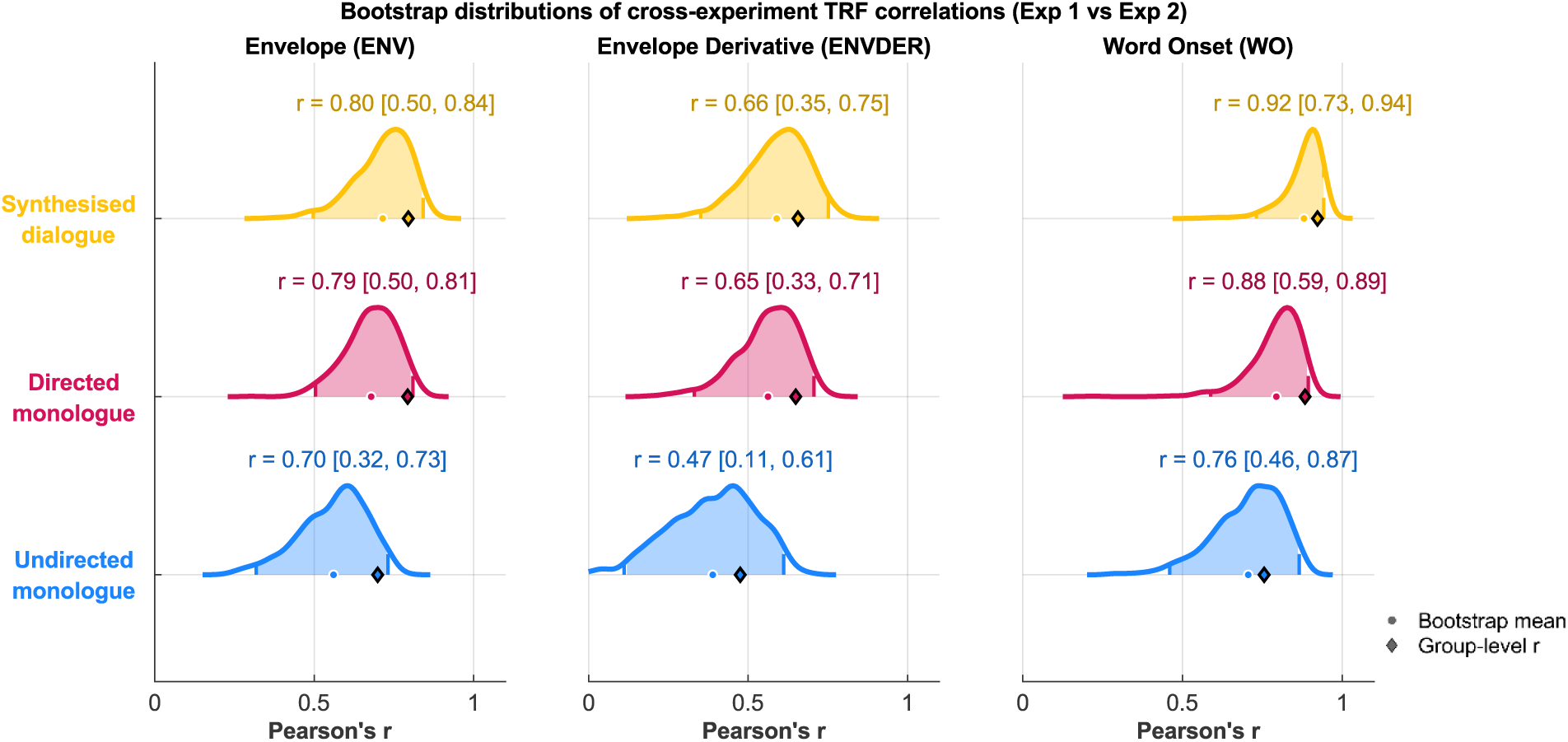
Speech podcast TRFs are more correlated with synthesised speech TRFs involving socially-relevant speech than undirected monologues. Correlations between group-level average TRF weights of the 64 EEG channels were calculated on the original samples from the two experiments. A bootstrap resampling procedure was performed to assess the robustness of this pattern. The distribution and confidence intervals of the bootstrap, the bootstrap mean (dot), and the group-level average (diamond) TRF correlation are plotted. TRF weights for the real podcast dialogues in Experiment 2 shows high correlations with the three conditions in Experiment 1 in the following descending order: synthesised dialogue, directed monologue, undirected monologue. This pattern emerges for all speech features in the analysis, with statistically significant differences emerging only for the word onset features across all conditions.

**Figure 4.**
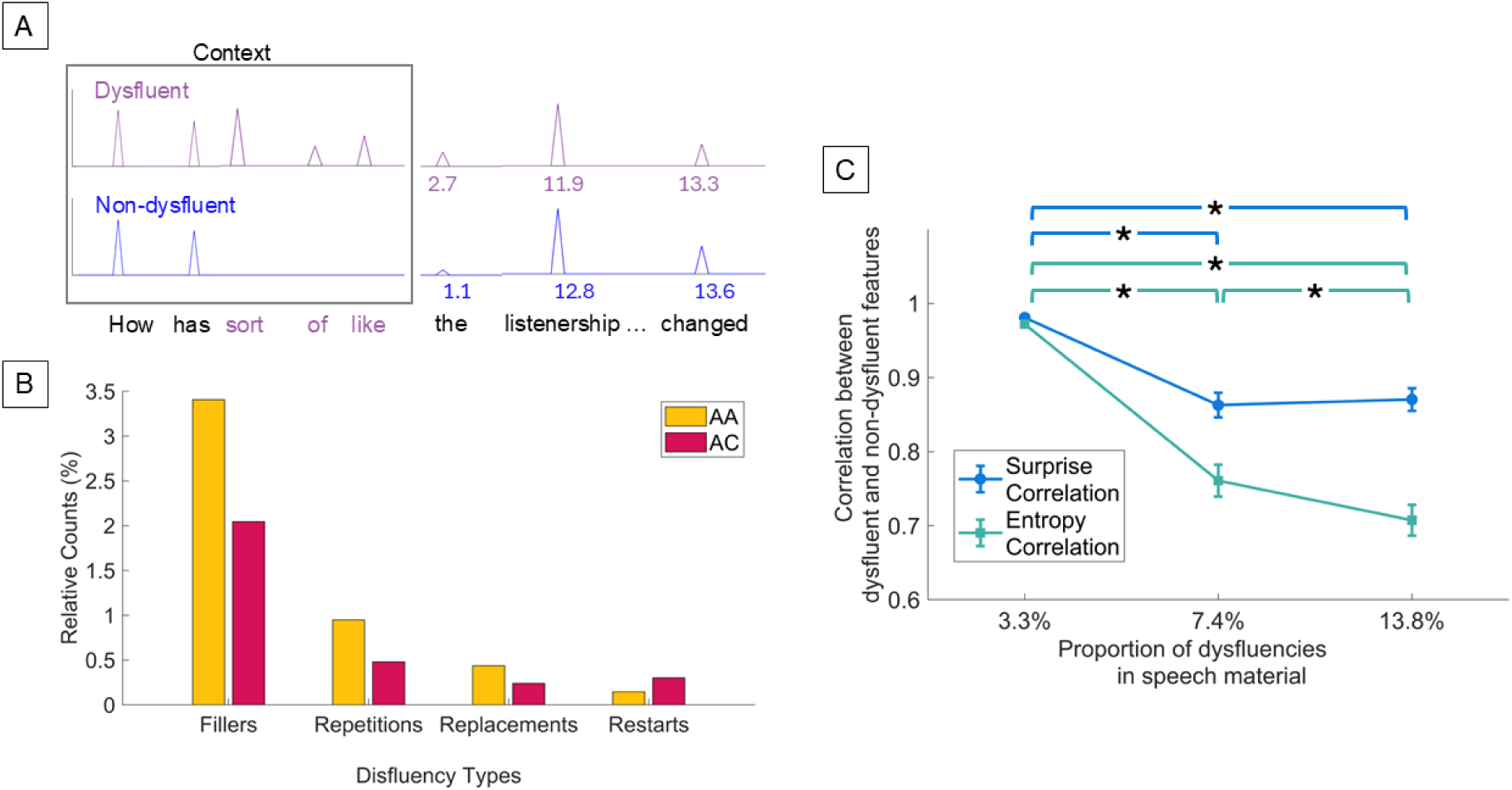
Investigating the impact of dysfluencies on LLMs. **(A)** Audio transcripts were analysed using two different lexical surprise and entropy vectors. While both vectors only included non-dysfluent words, for sake of comparison, lexical surprise and entropy were calculated using Mistral by including or excluding dysfluent words from the text input, producing the dysfluent and non-dysfluent feature vectors. **(B)** Dysfluencies were identified manually in the text material. We report the dysfluency frequency organised according to dysfluency type and experimental condition (adult-adult podcast AA; adult-child podcast AC). **(C)** The correlations between Mistral’s expectations when including or excluding dysfluent words in the calculation of lexical surprise and entropy are shown, quantifying the potential impact of dysfluency on the measurement of lexical expectations. Doubling the number of dysfluencies led to a statistically significant effect of dysfluency. On this basis, future research should measure dysfluencies in the speech material, since differences across participant cohorts or conditions may reflect differences in the prevalence of dysfluent words in the heard or produced speech.

To assess whether this pattern was statistically reliable across participants, conducted Friedman tests (due to Mauchly’s violation of sphericity), followed by post-hoc pairwise Wilcoxon signed-rank tests by considering each participant as a datapoint (N = 20). This analysis revealed a statistically significant effect of condition on the inter-experiment correlation when considering the word onset TRFs (χ²(2) = 11.70, *p* = 0.003, Kendall’s *W* = 0.29), while no statistical significant differences were detected for envelope (χ²(2) = 2.50, *p* = 0.29), and envelope derivative TRFs (χ²(2) = 4.90, *p* = 0.09). Post-hoc tests on word onset TRFs indicated that synthesised dialogue TRFs were significantly more correlated with real dialogue TRFs from Experiment 2 than directed monologue TRFs (Wilcoxon signed-rank *r*_synth-Dialog vs. real-Dialog_ > *r*_directed-M vs. real-Dialog_: *p* < 0.001), with the correlation further decreasing for monologue TRFs (*r*_directed-M vs. real-Dialog_ > *r*_undirected-M vs. real-Dialog_: *p* < 0.001) from Experiment 1. This trend is consistent with the correlation pattern synthesised dialogue > directed monologue > undirected monologue as on the original sample.

Additional analyses were carried out to obtain a more detailed assessment of the distribution and robustness of the cross-experiment correlations. Specifically, we performed a bootstrap resampling procedure (with replacement) and generated 1,000 correlations between the group-averaged TRF in Experiment 2 to each of the conditions in Experiment 1 for each feature. Compatibly with the participant-level results, the bootstrap distributions indicate particularly robust inter-experiment correlations for word onset TRFs, especially for synthesised dialogue.

#### Investigating the impact of dysfluencies on lexical expectation measurements

Experiment 2 involved real podcast recordings, which include dysfluencies by nature. Indeed, the dysfluency types and frequencies can vary due to a variety of factors (e.g., speaker, social context, social anxiety, fatigue, familiarity with interlocutor, knowledge of topic, speaking a native or non-native language). To investigate the impact of dysfluencies on the lexical surprise and entropy feature and its corresponding brain response, two lexical surprise and entropy vectors were constructed, including and excluding dysfluencies in the speech material, respectively (**Figure 4A**). As such, we carried out a characterisation of the dysfluency type and frequency for this specific dataset split across conditions AA and AC (**Figure 4B**), finding a majority of filler-type dysfluency and an overall frequency of dysfluent words of 3.36%, whereby ∼5% of all words were filler-type dysfluencies (e.g., “uh”/”like”/”you know”), ∼1% were repetitions, with less than 1% consisting of replacements and restarts (see **Materials and Methods** and **Table S1**). The frequency distribution of these dysfluencies types was consistent across podcast episodes, with a statistically significant main effect of dysfluency type (repeated measures ANOVA: *F*(4,60) = 28.49, *p* < 0.001, η² = 0.42; *post hoc* tests in **Table S2**). We found no statistically significant interaction of dysfluency type and experimental condition (adult-adult podcast AA; adult-child podcast AC) (repeated measures ANOVA: *F*(4,60) = 1.04, *p* = 0.39, η² = 0.02).

Here, we investigated if the presence of dysfluent words in the speech material (and its transcript) significantly impacts our ability to model the neural encoding of lexical expectations. To that end, we introduced two additional stimulus features in Experiment 2. These comprised word onset vectors modulated by lexical surprise and entropy values calculated using Mistral (see **Materials and Methods**). The first vector involved surprise values calculated when including all words, of which 3.3% of the were dysfluent. The second vector only included non-dysfluent words (dysfluencies were identified manually). However, dysfluent and non-dysfluent surprise values did not differ in this dataset, with a Pearson’s correlation between the two surprise vectors of 0.98. Due to this high correlation, we also did not expect differences to report in the coupling between these vectors and the corresponding EEG signals. While we could measure a statistically significant neural encoding of lexical surprise (using either dysfluent or non-dysfluent vectors) when compared to a baseline acoustic model (**Figure S3**), this is not guaranteed to be the case in speech material with more dysfluencies.

These virtually identical regressors suggest that at this low volume of dysfluencies, using speech material that includes or excludes dysfluencies would render little to no difference in the neural processing and encoding of speech. However, social speech can widely vary in the number of dysfluencies depending on numerous factors (e.g., speaker, social context, anxiety, tiredness, familiarity with interlocutor, knowledge of topic, speaking a native or non-native language). As a consequence, the neural encoding of speech may be impacted when more dysfluencies are present. An excessive number of dysfluencies (e.g., more dysfluent than non-dysfluent words) may disrupt lexical prediction in both the LLM and the human brain. Hence, we then assessed what level of dysfluency frequencies constitute an issue for lexical prediction in LLMs and to what extent this sensitivity persists by manually increasing the number of dysfluencies in the speech transcript. Specifically, we doubled and quadrupled the existing dysfluencies in Experiment 2, calculating the correlation between the lexical surprise and entropy values between the dysfluent and non-dysfluent vectors (**Figure 4C**). Lower correlations indicate a stronger impact of dysfluencies. We found a significant impact of dysfluency frequency (FDR-corrected Wilcoxon paired sign-ranked test: *p_originalSur vs. doubledSur_* < 0.001*, p_originalSur vs. quadSur_* < 0.001, *p_originalEntro vs. doubledEntro_* < 0.001, *p_doubledEntro vs. quadEntro_* < 0.001, *p_originalEntro vs. quadEntro_* < 0.001), whereby there was practically no sensitivity to dysfluencies for the frequency in the original material (dysfluent-nondysfluent correlation ∼ 1) and significant sensitivity to dysfluencies once they were doubled, though this effect seems to level out. Additionally, the strongest sensitivity overall emerged for entropy vectors (correlation down to ∼0.7).

## Discussion

Speech communication is an ideal testbed for investigating human cognition, as it involves a multitude of processes involved in the transformation of sound to meaning. Such processes encompass mechanisms such as statistical prediction, attention and cognitive effort. The possibility of extending that work to social speech opens up opportunities for probing a whole range of processes that are typically excluded from monologue listening research, such as social role and speaker familiarity, communication accommodation, social anxiety, and many others (Di Liberto & Ip, 2025). However, this opportunity comes with the important challenge of having to isolate the phenomena of interests from a much richer and harder-to-control set of behaviours and neural processes. One key for disentangling those processes and phenomena is to pinpoint how exactly these socially-relevant factors impact the spatial and temporal patterns reflecting the neural processing of speech and language. Here, we take a first step in that direction by means of a principled step-by-step approach, determining the impact of including a social element in speech listening tasks (excluding interaction in our scope of investigation).

The results from Experiment 1 indicate that social relevance substantially enhances the neural tracking of speech. Importantly, this effect cannot be explained by differences in low-level acoustic or speech-structure properties across stimulus categories. To assess this possibility, we quantified several stimulus characteristics, including duration, root-mean-square (RMS) amplitude, mean fundamental frequency (F0), F0 variability, and word rate. No systematic differences were observed between conditions, nor between monologue and dialogue stimuli. These findings indicate that the observed neural effects are unlikely to be driven by basic acoustic or speech-rate differences and instead reflect modulation by socially relevant aspects of the stimuli. The increased encoding emerged specifically for the speech envelope, but not for word-level processing (EEG prediction correlation metric; **Figure 1A**). Previous work on selective attention has also shown similar increases in envelope tracking.

Interestingly, however, the increased speech tracking in selective attention tasks also corresponded with changes in the TRF patterns, especially with a modulation of the TRF-P2 component (Carta et al., 2024; Power et al., 2012; Vanthornhout et al., 2018), while the increased envelope tracking here does not correspond to changes in the temporal or spatial dynamics of the TRF (**Figure 1D**). As such, our results are compatible with a general increased engagement with the task and stimulus, but with important differences from the widely studied selective attention phenomenon. It should be noted that we also detected sensitivity to the social relevance of speech in the beta band, specifically for word onset encoding. This result supports the potential and relevance of measuring cortical tracking at higher EEG rates (even though stronger speech tracking typically emerges in the delta and theta bands). While we did not have specific hypotheses pertaining to the EEG beta band in this study, we note that studying speech tracking at different frequency rates may provide an additional dimension for more clearly disentangling the cortical tracking of distinct speech features, and should be considered (without forgetting about the risks that come with filtering operations) (De Cheveigné & Nelken, 2019).

While Experiment 1 isolated the impact of social relevance on speech tracking, our design in Experiment 2 takes us a step further by measuring neural dynamics in more naturalistic settings, wherein spontaneous speech introduces additional complexities. The results in Experiment 2 (**Figure 2**) demonstrated temporal response characterisations that are not only consistent with previous monologue-listening literature (Broderick et al., 2018; Di Liberto et al., 2021; Heilbron et al., 2022), but also with the neural speech encoding results obtained in Experiment 1 (**Figure 3**). Our results indicated similar spatial and temporal patterns for synthesised and real speech recordings; the EEG encoding of speech features, including the speech envelope and word onset, was highly correlated across synthesised and real speech recordings, achieving correlations of up to 0.92 for word onset TRFs. This is quite remarkable, considering that the two experiments involved completely different speech material and distinct participant cohorts. The fact that we observed a consistent increase in that similarity with social relevance is also a strong indication that the impact of social relevance on the spatial and temporal patterns of speech is consistent across material and participants. Further work is needed to better characterise and test the boundaries of consistency in neural tracking of speech features across synthesised and real speech and to characterise the sensitivity of TRFs to variations in synthesiser model and configuration. Nonetheless, our findings suggest that the similarity between synthesised and real speech TRFs and their modulation by the social relevance of speech are remarkably high.

Experiment 2 demonstrates the feasibility of employing the TRF using naturalistic and socially-relevant speech material. Our dysfluency analysis indicated that the presence of dysfluencies observed in the podcast stimuli utilised does not impact the ability to derive indices of surprise and entropy, which can, in turn, be used to improve TRF models. This is promising for future research, as it suggests the possibility of studying speech tracking even in presence of dysfluencies. However, note here that the LLM dysfluency simulation analysis highlights that when a larger portion of the speech material includes dysfluencies, they can exert a notable impact on lexical expectation indices.

From this analysis, two key conclusions emerge. First, dysfluencies do not pose a problem up to a certain threshold, which may eliminate the need for manual transcript cleaning when using LLMs. Second, it is essential to quantify and report the dysfluency profile - both type and frequency - used in a given experiment, particularly when generating new stimuli. This task is relatively straightforward in listening experiments, where all participants are exposed to identical stimuli. In contrast, speech interaction paradigms, such as hyperscanning studies, introduce greater methodological complexity. A central challenge is that individual participants often exhibit unique dysfluency profiles (Befi-Lopes et al., 2014; Pistono et al., 2024), which can differentially affect measures of lexical expectations - both in terms of LLM-based estimation and neural encoding (Goldstein et al., 2022). This variability reinforces the need for systematic dysfluency measurement and individualised reporting to ensure robust and interpretable findings. Finally, it is worth noting that the observed sensitivity of LLMs to dysfluency opens new avenues for research focused on its impact on the neural encoding of speech. Future investigations could specifically examine this effect in participant cohorts with speech production challenges, such as individuals who stutter or those experiencing social anxiety. Such studies would not only deepen our understanding of speech processing under naturalistic conditions but also offer insights into the neural mechanisms underlying communicative difficulties.

In summary, this study demonstrates that incorporating social relevance into speech listening tasks enhances the neural tracking of speech and reveals remarkable consistent spatial-temporal patterns across varied speech materials and participant cohorts. By extending analysis to spontaneous speech and dysfluency, we demonstrate the possibility of modelling the neural tracking of speech, as well as lexical prediction mechanisms even in presence of dysfluencies. Based on our simulation, we provide evidence for the importance of considering the issue of dysfluency when studying social speech, when listening to speech interactions and especially in experiments where participants engage in dialogue. Altogether, these findings lay the groundwork for future investigations into the neural encoding of socially complex and dysfluent speech, offering a principled path toward understanding communication in real-world contexts.

## Supplementary

**Figure S1.**
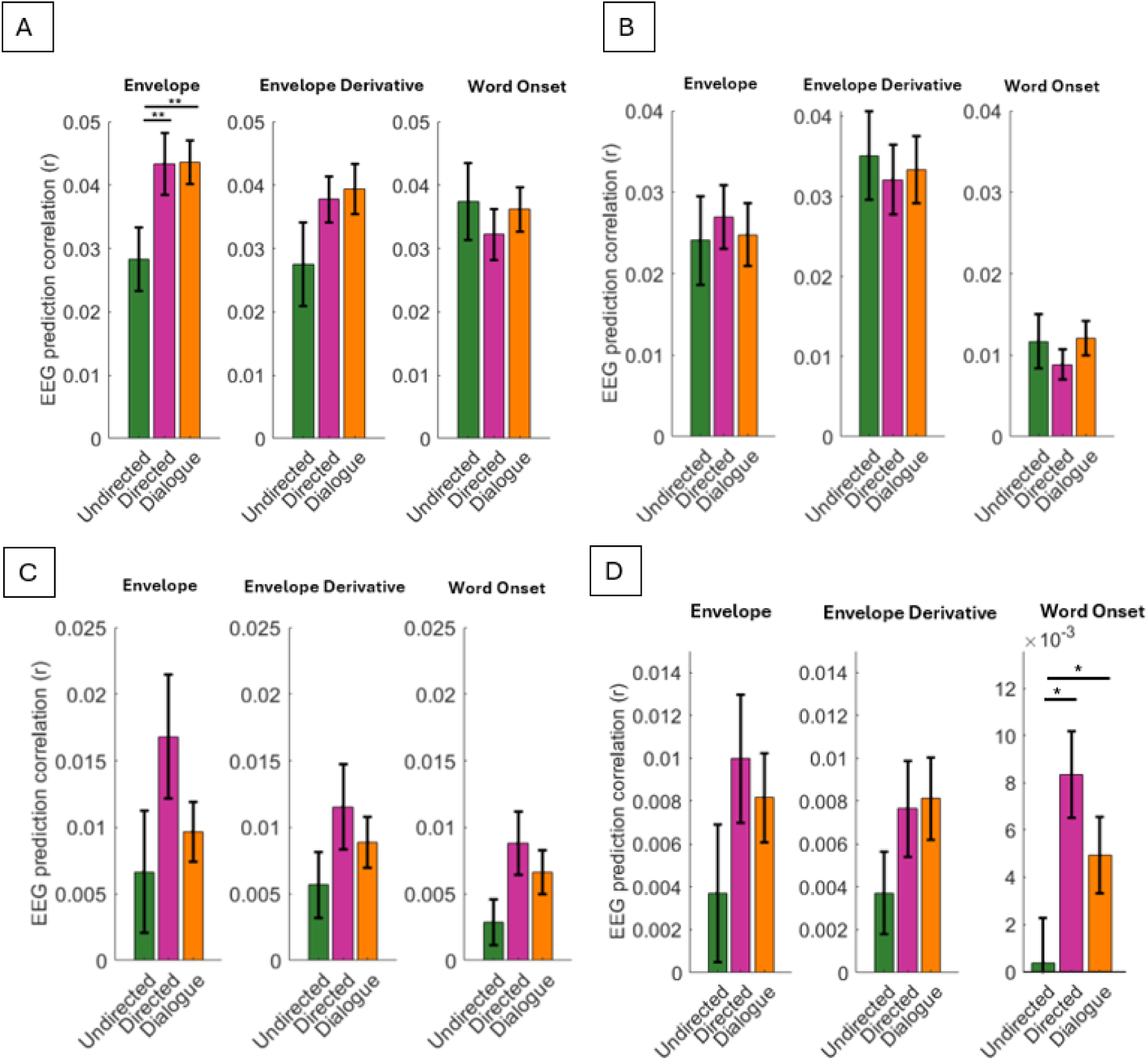
EEG prediction correlations for different frequency bands (Experiment 1). Univariate forward TRFs were fitted for envelope, envelope derivative, and word onset features to predict the EEG signal at frequency bands delta (A), theta (B), alpha (C), and beta (D). EEG prediction correlations (Pearson’s r), averaged over all channels and participants, are presented as mean ± SEM, comparing the three experimental conditions: undirected monologue, directed monologue, and dialogue. One-way ANOVA and Wilcoxon signed-rank post hoc tests were conducted (*p < 0.05, **p < 0.01). This data extends the results in Figure 1.

**Figure S2.**
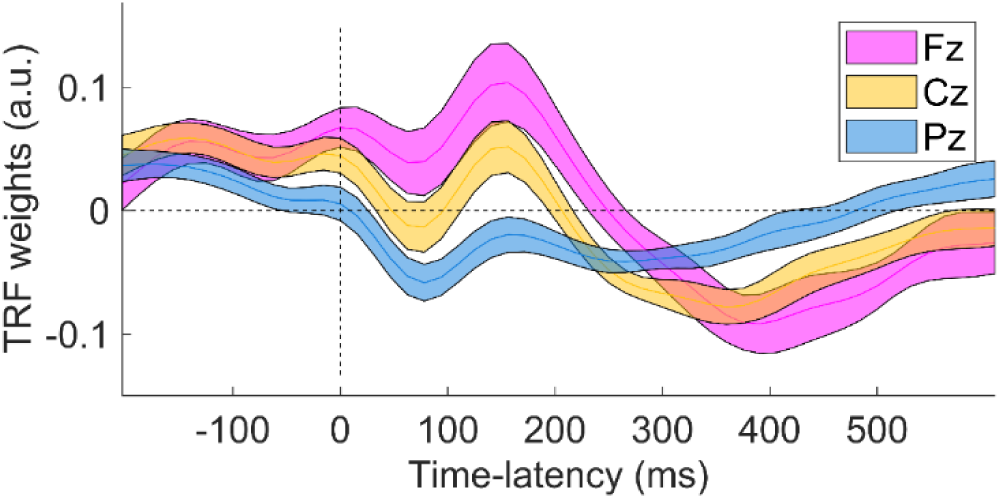
TRF weights for the envelope derivative feature from Experiment 2 at three selected EEG channels. This data extends the results in Figure 2.

**Table S1.** Dysfluency types. The dysfluency analyses in Figures 4 and 5 constitute a preliminary approach for handling dysfluencies that are a natural part of social speech. This table indicates the four dysfluency types that were identified – fillers, repetitions, restarts (also known as false starts), and replacements (also known as repairs) (Shriberg, 1999). See Figure 4B for the distribution of these dysfluency types in the speech material for Experiment 2. Note that Experiment 1 did not include any dysfluencies.

| Dysfluency Type | Definition |
| --- | --- |
| Fillers | uh / um, you know, sort of, like, I'll say, I don't know |
| Repetitions | I-I-I |
| Restarts (also coined *false start* in (Shriberg, 1999)) | *The speaker abandons completely the initial utterance and restarts it. This type of dysfluency does not actually involve a repair as the speaker begins a completely new utterance*(Passali et al., 2022). |
| Replacements (also coined *repair* in (Shriberg, 1999)) | *The speaker replaces the dysfluent word(s) or phrase with the fluent one. In replacements, the reparandum is replaced by the repair* (Passali et al., 2022) |

**Table S2.**
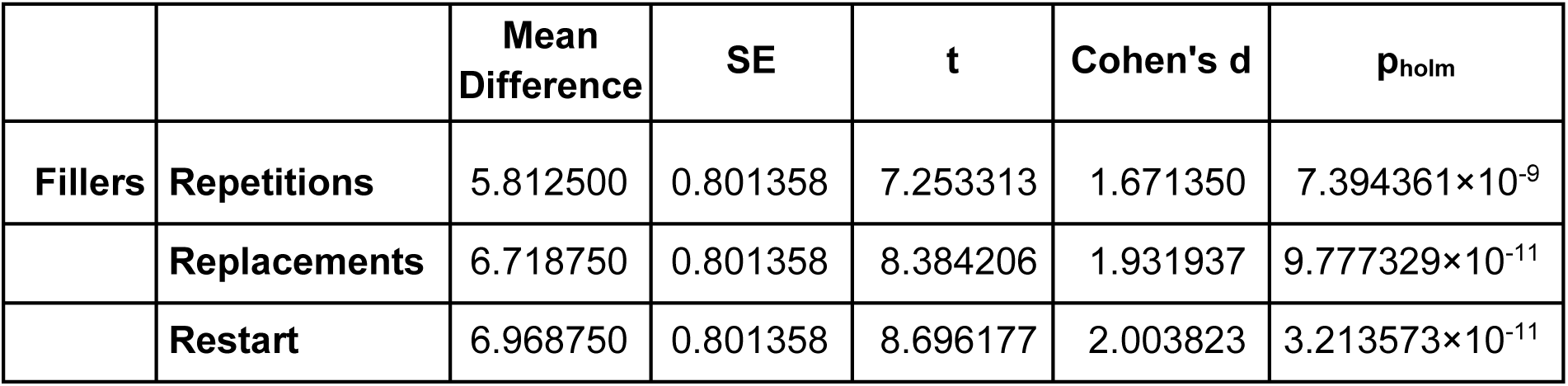
Post Hoc Comparisons of dysfluency types. A statistically significant main effect of dysfluency type emerged when comparing the frequency distribution of dysfluencies across trials in both AA and AC conditions with a repeated measures ANOVA. Post hoc tests show how fillers significantly outnumber any other dysfluency produced in real podcast dialogues. This data extends the results in Figure 4B.

**Figure S3.**
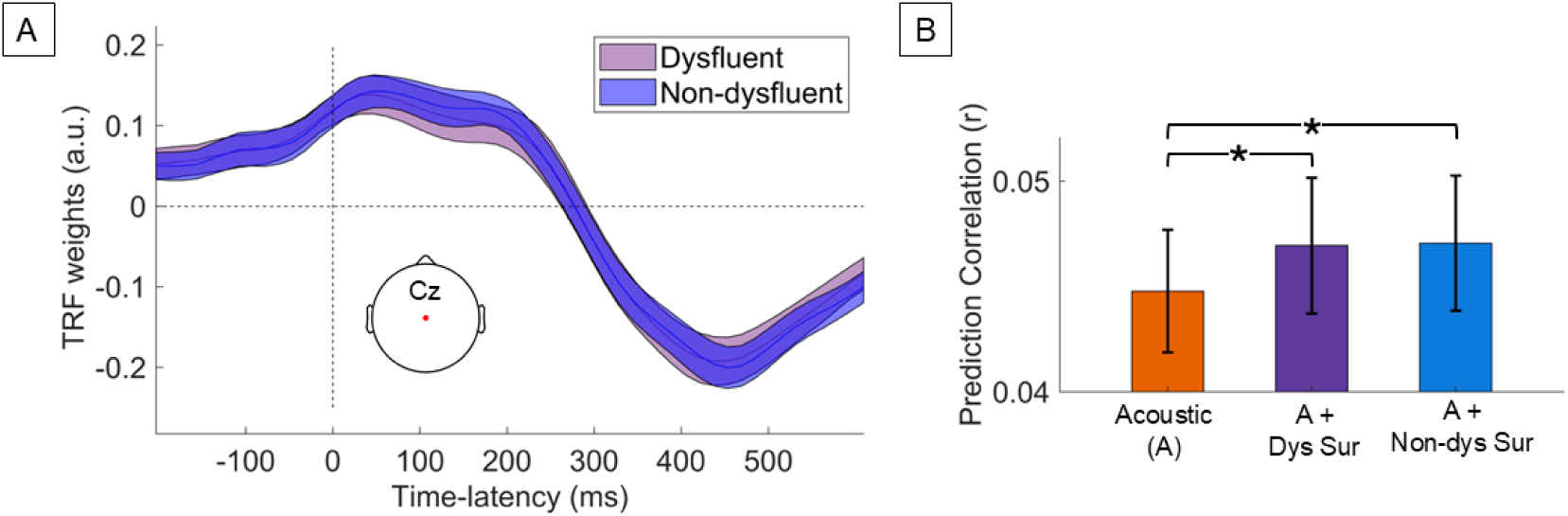
Investigating the impact of dysfluencies on speech TRFs. **(C)** Multivariate TRFs were computed to predict the EEG signal using the dysfluent or non-dysfluent vectors, concatenated with nuisance regressors capturing speech acoustics and word segmentation (envelope, envelope derivative, and word onset). We report the TRF weights corresponding to the dysfluent and non-dysfluent surprise features at electrode Cz. **(D)** Both versions of the models that included acoustics and dysfluent surprises returned a significant gain in the average EEG prediction correlations in comparison to the acoustics-only mTRF, reflecting a statistically significant neural encoding of lexical surprise. These data extend the results in Figure 4.

Forward multivariate TRFs were fit for each of the two vectors and including the three features used in the earlier analysis as nuisance regressors (envelope, envelope derivative, and word onsets), leading to a *non-dysfluent surprise TRF* and a *dysfluent surprise TRF*. Note that, for sake of comparison, dysfluent and non-dysfluent vectors were built to be identical in everything but the surprise values i.e., they had the same word onsets. Finally, a third TRF was fit for the nuisance regressors without any lexical surprise information (*baseline* TRF). When comparing the model weights between the non-dysfluent and dysfluent surprise TRFs (Wilcoxon paired signed-rank test: *p* = 0.951; **Figure S3a**), no statistically significant differences emerged. In addition, EEG prediction correlations were compared across the three TRF models, showing a statistically significant differences across the three TRF models (repeated measures ANOVA: *F*(1, 19) = 7.96, *p* = 0.001, η² = 0.30; **Figure S3b**). *Post hoc* comparisons indicated larger EEG prediction correlations when including either surprise vector (*t*-tests; *r*_non-dysfluent surprise_ > *r*_no-surprise_, *t*(19) = -3.54, *p_holm_* = 0.003, Cohen’s *d* = -0.16; *r*_dysfluent surprise_ > *r*_no-surprise_, *t*(19) = -3.36, *p_holm_* = 0.004, Cohen’s *d* = -0.16). As expected, no statistically significant differences were measured between non-dysfluent and dysfluent surprise TRF prediction correlations (*t-*test: *t*(19) = -0.18, *p_holm_* = 0.861, Cohen’s *d* = -0.01).

**Figure S4.**
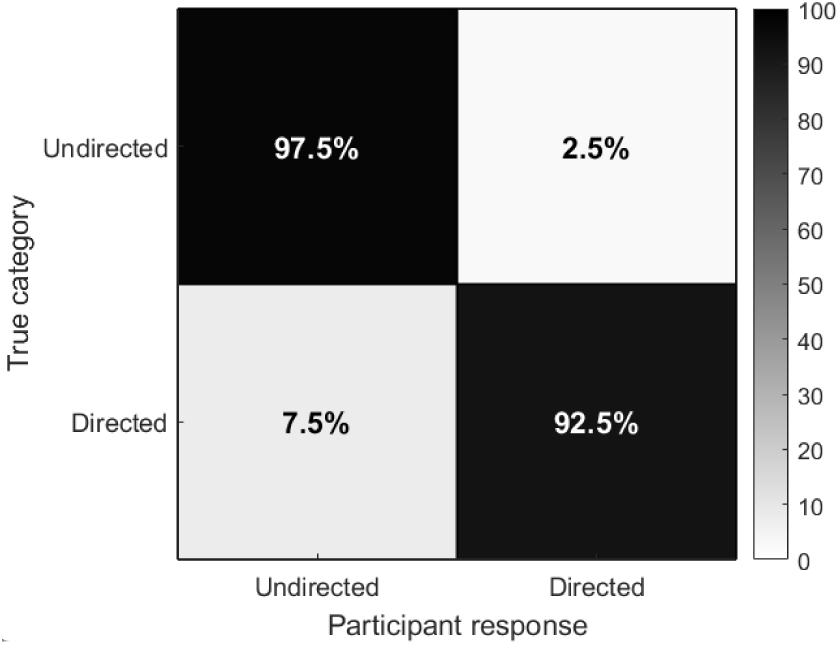
Behavioural validation of stimulus categorisation into directed vs. undirected. An independent group of participants (N = 15) was asked to categorise speech recordings as directed or undirected. 16 recordings, each ∼2-minute long, were used in this experiment. The confusion matrix reports correct and incorrect classification rates.

**Table S3.** Acoustic and speech-structure properties across conditions. No significant differences were observed between undirected monologue, directed monologue, and dialogue conditions, nor between monologue and dialogue stimuli.

| Measure | Comparison | Test | *p*-value | Effect size |
| --- | --- | --- | --- | --- |
| Duration | Undirected vs Directed vs Dialogue | Kruskal–Wallis | 0.57 | 0.01 ($\epsilon^2$) |
| RMS amplitude | Undirected vs Directed vs Dialogue | Kruskal–Wallis | 0.17 | 0.08 ($\epsilon^2$) |
| Mean F0 | Undirected vs Directed vs Dialogue | Kruskal–Wallis | 0.93 | 0.01 ($\epsilon^2$) |
| F0 variability | Undirected vs Directed vs Dialogue | Kruskal–Wallis | 0.40 | 0.02 ($\epsilon^2$) |
| Word rate | Undirected vs Directed vs Dialogue | Kruskal–Wallis | 0.15 | 0.09 ($\epsilon^2$) |
| Duration | Monologue vs Dialogue | Wilcoxon rank-sum | 0.30 | 0.18 (r) |
| RMS amplitude | Monologue vs Dialogue | Wilcoxon rank-sum | 0.51 | 0.12 (r) |
| Mean F0 | Monologue vs Dialogue | Wilcoxon rank-sum | 0.72 | 0.06 (r) |
| F0 variability | Monologue vs Dialogue | Wilcoxon rank-sum | 0.51 | 0.11 (r) |
| Word rate | Monologue vs Dialogue | Wilcoxon rank-sum | 0.32 | 0.18 (r) |

## Acknowledgements

This work was conducted with the financial support of the Research Ireland Centre for Research Training in Digitally-Enhanced Reality (d-real) under Grant No. 18/CRT/6224. This research was conducted with the financial support of Research Ireland at ADAPT, the Research Ireland Centre for AI-Driven Digital Content Technology at Trinity College Dublin and University College Dublin [13/RC/2106_P2]. For the purpose of Open Access, the authors have applied a CC BY public copyright licence to any Author Accepted Manuscript version arising from this submission.

## Footnotes

1 https://app.play.ht/

2 https://osf.io/hx3nz

3 https://laist.com/podcasts/servant-of-pod/kids-podcasts-a-true-alternative-to-screen-time;https://creators.spotify.com/pod/profile/martine-severin/episodes/31--How-to-Become-an-Epic-Improv-Comic-with-Joy-Dolo-e1js9c2

4 https://github.com/CNSP-Workshop/CNSP-resources/tree/main/CNSP/exampleScripts

5 https://cnspworkshop.net/

